# Siglec-15 is a glyco-immune checkpoint in prostate cancer regulating immune evasion and metastasis

**DOI:** 10.64898/2026.08.07.743480

**Authors:** Nicoll Matthews, Feier Zeng, Kirsty Hodgson, Matthew Fisher, Ziqian Peng, Libby Blencoe, Margarita Orozco-Moreno, Ella P Dennis, Liancheng Lu, Michelle A Lawson, Shenglin Mei, David B Sykes, Dallas B Flies, Richard Beatson, Ning Wang, Jennifer Munkley

## Abstract

Prostate cancer is a leading cause of cancer-related mortality in men, and effective treatment options are limited for advanced and metastatic disease. The sialoglycan immune checkpoint Siglec-15 has emerged as a key mediator of tumour-associated immune suppression in several malignancies; however, its expression and functional role in prostate cancer remain poorly defined. Here, using dual immunofluorescence and immunohistochemistry, we demonstrate that Siglec-15 is expressed by prostate tumour epithelial cells, immunosuppressive macrophage phenotypes, and bone-resorbing osteoclasts within the tumour microenvironment. Mechanistically, we show that direct Siglec-15 receptor crosslinking, either by antibodies or tumour cell-derived conditioned medium, promotes monocyte-to-macrophage differentiation, generating macrophages with immunosuppressive and pathogenic phenotypes. Using therapeutic antibodies, we show that Siglec-15 blockade suppresses supernatant-induced monocyte to macrophage differentiation, allowing for the recovery of CD8⁺ T-cell activation. Furthermore, we reveal that macrophage colony-stimulating factor (M-CSF) driven monocyte-derived macrophage differentiation is partially dependent on Siglec-15 signalling, with Siglec-15 blockade enhancing CD8⁺ T-cell responses. In addition, anti-Siglec-15 treatment suppressed osteoclast differentiation, highlighting a dual role for Siglec-15 in prostate cancer immune suppression and bone remodelling. Consistent with these *in vitro* findings, therapeutic Siglec-15 blockade significantly reduced subcutaneous tumour growth in a CD8⁺ T-cell-dependent manner and prolonged survival in a mouse model of prostate cancer metastasis. Together, these findings identify Siglec-15 as a central regulator of the prostate cancer glyco-immune axis, linking tumour-associated macrophage immune suppression with osteoclast-mediated bone remodelling, providing a compelling rationale for the clinical development of Siglec-15-targeted therapies for patients with advanced disease.

## INTRODUCTION

Prostate cancer is a major cause of cancer related mortality in males worldwide and remains challenging to treat once metastatic disease has developed (1). Although advances in androgen receptor (AR) targeted therapies have improved outcomes, most patients ultimately develop castration resistant prostate cancer (CRPC), with bone representing the dominant and most clinically significant site of metastasis (2). Bone metastasis can lead to substantial morbidity, including pain, fractures, and reduced quality of life, and there is an urgent unmet clinical need to identify new therapeutic approaches (3). Aberrant glycosylation is a hallmark of cancer and tumour-associated glycans play functional roles in disease progression (4, 5). Alterations to sialylated glycans (sialoglycans) are particularly prominent in cancer, where tumour-associated changes in sialylation contribute to disease progression (6, 7). Sialoglycans engage Siglec receptors expressed on immune cells, forming an inhibitory glyco-immune checkpoint that suppresses anti-tumour immune responses (8). Distinct from classical protein based immune checkpoints, the Siglec–sialoglycan axis plays a key role in regulating tumour immune interactions, and is a growing area of interest in cancer glycobiology (9, 10).

In prostate cancer, altered sialyltransferase expression and elevated tumour sialylation have been linked to tumour growth, metastasis, and immune evasion (11–16). Our previous work and that of others has demonstrated that prostate tumours exhibit increased levels of Siglec-engaging sialoglycans that can promote an immunosuppressive tumour microenvironment and have highlighted the clinically actionable nature of this axis in prostate cancer (16–19). While global desialylation strategies using potent metabolic inhibitors or sialidase enzymes have demonstrated efficacy to impede prostate cancer progression (15, 20, 21), the specific role of individual Siglec-sialoglycan interactions within this axis remains to be fully mapped. Siglec-15 is a central glyco-immune checkpoint, acting both to suppress T cell responses and modulate bone remodelling through osteoclast differentiation (22–26). Studies in breast cancer identify the Siglec-15 sialoglycan axis as a modulator of tumour-induced osteoclastogenesis that is targetable to suppress bone metastatic breast cancer (25), providing a strong basis to also investigate this pathway in prostate cancer. However, the mechanisms through which Siglec-15 coordinates monocyte fate, macrophage survival, and bone remodelling under the influence of the prostate tumour microenvironment are not yet understood.

Here, we investigated the expression and functional role of Siglec-15 in prostate cancer, with a focus on its contribution to immune evasion and metastasis. Using dual immunofluorescence and immunohistochemistry, we show Siglec-15 is expressed by prostate tumour epithelial cells, immunosuppressive macrophages, and osteoclasts within the prostate tumour immune microenvironment. Mechanistically, we demonstrate that direct Siglec-15 receptor crosslinking alone acts as an inductive switch to initiate monocyte differentiation, and that macrophage colony stimulating factor (M-CSF) can exploit this axis to program monocytes into an immunosuppressive state. Furthermore, using primary human immune cell systems, we show that therapeutic antibody blockade of Siglec-15 reduces immunosuppressive macrophages, or alters their phenotype, whilst decreasing their secretion of transforming growth factor beta (TGF-β). This in turn alleviates the suppression of CD3/CD28 mediated CD8⁺ T cell proliferation and interferon gamma (IFN-γ) release. Finally, we establish that therapeutic blockade of Siglec-15 impairs *in vivo* subcutaneous tumour growth in a CD8⁺ T-cell-dependent manner, inhibits tumour-induced osteoclastogenesis, and significantly prolongs survival in a mouse model of prostate cancer metastasis. Together, our findings identify the Siglec-15 sialoglycan axis as a dual action therapeutic vulnerability, linking myeloid mediated immune suppression with bone remodelling, and providing rationale for the clinical development of Siglec-15-targeted therapies for patients with advanced prostate cancer.

## METHODS

### Immunofluorescence (clinical tissue)

FFPE-fixed prostate clinical tissues were dewaxed and rehydrated before antigen retrieval at 95°C for 20 min using Sodium Citrate Antigen Retrieval Buffer (Proteintech, PR30001). Tissues were then blocked with 1X CFB (Vector Laboratories, SP-5040-125) for 1 hr at room temperature, followed by overnight incubation in the dark at 4°C in 1:100 Siglec-15 (Abcam, AB198684), 1:200 CD206 (Proteintech, 60143-1-1G), 1:200 Cathepsin K (Santa Cruz, sc-48353), 1:500 AMACR/p504S Rabbit Polyclonal Antibody (Proteintech, 15918-1-AP), 1:200 AMACR Mouse Monoclonal Antibody [UMAB68] (UltraMAB, UM870012), or 10ug/mL rh Siglec-15 Chimera (R&D systems, 9227-SL). Next, tissues were washed with PBS before 1 hr incubation in the dark at room temperature in 1:1000 Multi-rAb CoraLite Plus 594 Goat Anti Rabbit Recombinant Secondary Antibody (Proteintech, RGAR004), 1:500 Goat Anti-Mouse IgG H & L Alexa Fluor 488 (Abcam, ab150113) or 1:500 Nano-Secondary Alpaca Anti-Human IgG recombinant VHH CoraLite Plus 647 (Proteintech, CTK0117). Finally, following PBS washes, tissues were mounted with VectaShield AntiFade Mounting Medium with DAPI (Vector Laboratories, H-1800). Images were acquired and processed with the ZEISS Axio Imager 4 and ZEN Microscopy Software (blue edition).

### Clinical cohorts

#### TMA cohort 1

A previously published 40 case human prostate cancer TMA (20) was purchased from Novus Bio (NBP2-30169). This TMA includes 4 µm thick, 2.0 mm diameter single cores per case.

#### TMA cohort 2

A previously published 96 case TMA (19) comprising cores of normal prostate tissue and prostate adenocarcinoma of different Gleason grades was purchased from US Biomax (PR1921-L38). For this TMA, there are 5 µm thick, 1.5 mm diameter duplicate cores per case. The TMA is an updated version of US Biomax TMA PR1921b, which has been previously published by us (27, 28).

#### TMA cohort 3

A previously published TMA (29) containing matched normal prostate tissues and prostate tumour tissues from 60 patients. This TMA includes 5 µm thick, 1.5 mm diameter triplicate cores per case, and was kindly gifted to us by Professor Colm Morrissey (University of Washington).

#### TMA cohort 4

100 case CRPC metastatic clinical heterogeneity TMA intended to assess the heterogeneity of molecular markers across different anatomic sites in lethal prostate cancer metastasis (obtained via rapid autopsy). This TMA has been previously published by us (20) and was created as part of the Movember Global Action Plan 1 Unique tissue microarray (GAP1-UTMA) project (30).

#### LuCaP Patient-derived Xenograft (PDX) TMA

The LuCaP PDX TMA contains samples from previously published PDX model tumours (31, 32) implanted and grown subcutaneously (in triplicate) in intact or castrated mice. PDX tissues were established from distinct patients using specimens acquired at either radical prostatectomy or at autopsy. Castration resistant derivatives of each PDX model were generated by enabling the tumours to grow and progress in castrated mice. Cores were 5 µm thick,1.5 mm-diameter, and included in triplicate per LuCaP line. This TMA was kindly provided to us by Professor Eva Corey and Professor Colm Morrissey (University of Washington).

#### Primary prostate cancer tissue

Patient tissue samples were collected with ethical permission from Castle Hill Hospital (Cottingham, Hull) (Ethics Number: 07/H1304/121). Use of patient tissue was approved by the Local Research Ethics Committees. Patients gave informed consent and all patient samples were anonymised.

#### Bone metastasis tissue samples from rapid autopsy

20 cases of rapid autopsy FFPE tissue samples from prostate-derived tumours growing in bone were kindly provided by Dr Colm Morrissey (University of Washington) via the Prostate Cancer Biorepository Network (PCBN). Biopsies of metastatic bone sites were obtained from patients with CRPC within hours of death using a cordless drilling trephine (DeWalt Industrial Tool) and model 2422-51-000 trephine (DePuy). Bone cores were fixed in 10% neutral buffered formalin, decalcified with 10% formic acid and paraffin embedded.

### Immunohistochemistry

Immunohistochemistry to detect Siglec-15 protein in FFPE tissue sections was performed as described previously (14), using a well validated Siglec-15 antibody (clone 1F7) developed as a potential companion diagnostic for clinical trials (33) (provided by NextCure). Antigen retrieval was performed by heating to 95°C in a water bath for 20 minutes followed by staining the tissues with anti-Siglec-15 antibody (Siglec-15, clone 1F7, NextCure) (33) at a 1:2000 dilution. Nuclei were counterstained with haematoxylin. The TMAs were scored by a pathologist using the 0-300 Histoscore score method (34). Only epithelial cells were scored. Sections were scored based on their staining intensity with 0 being assigned to cells with absent staining, 1 to weak staining, 2 to moderate staining and 3 to strong staining. Within each staining intensity the percentage of epithelial cells (0–100%) with this staining intensity was assigned. This resulted in a Histoscore calculated from the following equation H = 0x (% of cells scored at 0) + 1x (% of cells scored at 1) + 2x (% of cells scored at 2) + 3x (% of cells scored at 3). For the LuCaP PDX TMA shown in Figure 3B, images were analysed using QuPath (version 0.6) software as described previously (19). Each PDX sample was analysed in triplicate samples from the same mouse, and across three independent mice. Positive cells were outlined and classified as having low, moderate or high staining intensity to calculate a 0-300 HistoScore, calculated as the sum of each staining intensity score (1+, 2+, 3+) multiplied by the percentage of cells classified at that intensity using the equation staining Index (H-score) = Σ (each intensity score × % of cells at that intensity). In line with previous studies (34), only epithelial tumour cells were scored.

### Siglec-15 therapeutic antibodies

Siglec-15 therapeutic antibodies were provided by NextCure for use in our study, including the anti-mouse Siglec-15 monoclonal antibody NP154 (Clone 5G12.mIgG2a Fc silenced), which is a surrogate of the parental antibody of the humanised clinical-stage anti-human Siglec-15 antibody NC318 and a respective antigen-binding deficient mutant control antibody NP154, Siglec-15 antibody (NP159 (Clone 1H3.mIgG2a Fc silenced) and a respective antigen-binding deficient mutant mIgG2a Fc silenced control antibody NP149.

### Generation of human monocyte-derived macrophages

Leucocyte cones were ordered from the National Health Service Blood and Transplant Service (NHSBTS). Cells were mixed 1:1 with phosphate-buffered saline (PBS) and layered on Ficoll–Paque (GE Healthcare, 1714402). Cells were spun at 400g for 30 min, with the brake off, and the human peripheral blood mononuclear cells (PBMCs) were taken from the buffy layer above the Ficoll–Paque. CD14+ cells were isolated from peripheral blood mononuclear cells using the MACS system (Miltenyi Biotech, 130-050-201, LS Columns, 130-042-401). CD14+ cells were plated at 1 × 10^6^/mL in 850 μL AIM-V media (ThermoFisher, 12055091) plus 150 μl concentrated supernatant from wild type CWR cells or 50ng/ml M-CSF (Biolegend, 574804) in the presence of 10ug/ml anti-Siglec-15 (NextCure, 5G12-mG2a) or mutant control (NextCure, 1129-mG2a). Supernatant, M-CSF and antibodies were replenished every 3 days. All cells were cultured for 6 days before harvesting.

### Stimulation of PBMCs

Anti-CD3 (Biolegend, 317325) was plated out at 1ug/ml in PBS on a 96 well plate and left overnight. PBMCs were isolated as above and stained with 5uM CFSE (Biolegend, 423801) as per the manufacturer’s instructions. The 96 well plate was washed with PBS and 100ul / test of supernatant from macrophages was taken from above and plated along with 1ug/ml anti-CD28 (Biolegend, 302933). 100ul of 1×10^6^/ml labelled PBMCs were added to each appropriate well and cells were incubated for 3 days. After 3 days, supernatant was collected for IFN-y measurement, and the cells were taken forward for flow cytometry.

### Flow cytometry (macrophages and PBMCs)

1 × 10^5^ cells were stained with a live/dead dye (ThermoFisher, L23102) in PBS for 10 min on ice in the dark, before being washed twice in FACS buffer (0.5% bovine serum albumin [Sigma; 05482] in PBS + 2 mM EDTA). PBMCs were then taken forward and measured for proliferation. Macrophages were Fc blocked with TruStain (Biolegend, 422302) in FACS buffer for 10 min on ice in the dark. Cells were washed and then stained with anti-CD206 (Biolegend, 321104), Anti-CD86 (Biolegend, 374206, anti-PD-L1 (Biolegend, 393606), anti-HLA-DR (Biolegend, 307628) or isotype controls (Biolegend, 400149, 400112, 400108), using concentrations recommended by the manufacturer, on ice for 30 min in the dark. Cells were then fixed, permeabilised and stained (anti-arginase; Biolegend, 369704 or anti-CD8a; Biolegend, 300908) as per manufacturer’s instructions (Biolegend, 421403). Cells were then washed and read using a BD Accuri C6 Plus flow cytometer, with analysis carried out using BD Accuri C6 Plus software. All cells were gated as follows: (a) Forward scatter and side scatter (SSC) to exclude cellular debris (whilst also adjusting threshold), (b) live/dead (only live cells carried forward) and (c) SSC-A vs. SSC-H—only singlets carried forward. MFIs were corrected against the isotype control. % PBMC proliferation was calculated against unstimulated cells +/- CD8+ gating.

### TGF-β and IFN-y ELISAs

Supernatant from indicated cells was assessed for TGF-β (1/2 dilution; R&D systems DY240) and IFN-y (1/100 dilution; Biolegend 430101) according to manufacturer’s instructions

### Osteoclast cultures and assays

Osteoclast formation was assessed in bone marrow (BM) and RAW264.7 cell cultures stimulated with receptor activator of nuclear factor κB ligand (RANKL, R&D, 462-TEC) and M-CSF (R&D, 416-ML-010), in the presence or absence of anti-Siglec15 antibody (10 μg/mL, provided by NextCure). BM cells were isolated from the long bones of 6–8-week-old mice and cultured in standard α-MEM supplemented with M-CSF (100 ng/mL) for 48 h to generate osteoclast precursors. M-CSF-dependent precursor cells were then seeded into 96-well plates (3 × 10 cells/well) and cultured with M-CSF (100 ng/mL) and RANKL (25 ng/mL) for 24 h. Mouse osteoclasts were also generated in cultures of mouse RAW 264.7 pre-osteoclast-like macrophages cultured in the presence of RANKL (100 ng/ml), prior to the addition of anti-Siglec-15 antibody (NC318, 10 mg/ml). Osteoclast cultures were terminated by fixation in 10% neutral buffered formalin, and stained for Tartrate-resistant acid phosphatase (TRAP). TRAP+ with 3 or more nuclei were considered to be osteoclasts and manually counted under a microscope.

### Mouse models

#### TRAMPC2 tumour allografts

Twelve 6-week old male C57BL/6 mice (RRID:MGI:2159769) were purchased from Charles Rivers and housed in a controlled environment. 2×10^6^ TRAMPC2 cells (35) resuspended in 100 μL (50% PBS+ 50% Matrigel) were injected subcutaneously into the left flank of mice. Mice were treated with 10 mg/kg control antibody NP154, or 10 mg/kg NP154 anti-Siglec-15 (Clone 5G12 IgG2aSi (Fc silenced) antibody or, NP154 anti-Siglec-15 antibody + anti-CD8 antibody twice a week via intraperitoneal injection. For *in vivo* depletion of CD8+ T cells, anti-CD8 depleting antibody (anti-mouse CD8a, 53-6.7, 2BScientific) was administered intraperitoneally at 10 mg/kg on days −2 and 0 relative to the time of tumour cell injection, followed by weekly treatments for the duration of the experiment. Tumour progression was monitored twice weekly using callipers. Mice were euthanised through exsanguination under general anaesthesia, followed by cervical dislocation.

### PC3 bone metastasis study

Six-week old male BALB/c nude mice (RRID:MGI:2683685) were purchased from Charles River (Kent, UK) and housed in a controlled environment in Optimice cages (Animal Care Systems, Colorado, USA) randomly located in the cage rack with a 12 hour light/dark cycle at 22°C with ad libitum water and 2018 Teklad Global 18% protein rodent diet containing 1.01% Calcium (Harlan Laboratories, UK). Twenty mice received single-cell suspensions of 1×10^5^ PC3 cells/100 μL PBS via injection into the left cardiac ventricle of mice (intracardiac injection). Mice were randomised into two groups to receive twice weekly dosing of either Siglec-15 antibody (NP159 Fc silenced, NextCure) or control antibody (NP149, NextCure) at 10 mg/kg via intraperitoneal injection starting at 2 days post tumour inoculation. Survival times were monitored over 6 weeks. Mice were monitored daily and euthanized upon reaching predefined humane endpoints, including, but not limited to, >15% body weight loss, poor body condition, or significant behavioural changes. The time to euthanasia was recorded and used to generate Kaplan–Meier survival curves. Mice were euthanized through exsanguination under general anaesthesia, followed by cervical dislocation. Procedures complied with the UK Animals (Scientific Procedures) Act 1986 and were reviewed and approved by the local Research Ethics Committees of the University of Sheffield under Home Office project licence PP3267943 (Sheffield, UK).

### Transcriptomic analyses

For the transcriptomic analysis in Figure 1 and Supplementary Figure 3, *SIGLEC15* expression was analyzed in three published single-cell RNA-sequencing datasets covering primary prostate cancer and prostate cancer bone metastasis. Primary and advanced tumour data were obtained from the prostate tumor meta-atlas of Zhang et al. (36). Bone metastatic prostate cancer datasets were downloaded from Kfoury et al. (GSE143791) (37) and Wang et al. (OEP005136) (38). We obtained the original Seurat object with cell annotation for the primary and advanced tumour datasets. For the bone-metastasis dataset, cells with fewer than 500 counts or more than 40% mitochondrial reads were removed and doublets were flagged per sample with Scrublet (39). Count matrices were normalized with log-transformation and reduced to 50 principal components on 2,000 highly variable genes. Batch effects were corrected with Harmony, and the corrected components were used for a k = 15 neighbor graph, UMAP and Leiden clustering, as implemented in Scanpy (40). Cell-type annotations were obtained from the original authors of each dataset and myeloid subclusters of the bone-metastasis datasets were named by maker genes.

**Figure 1.**
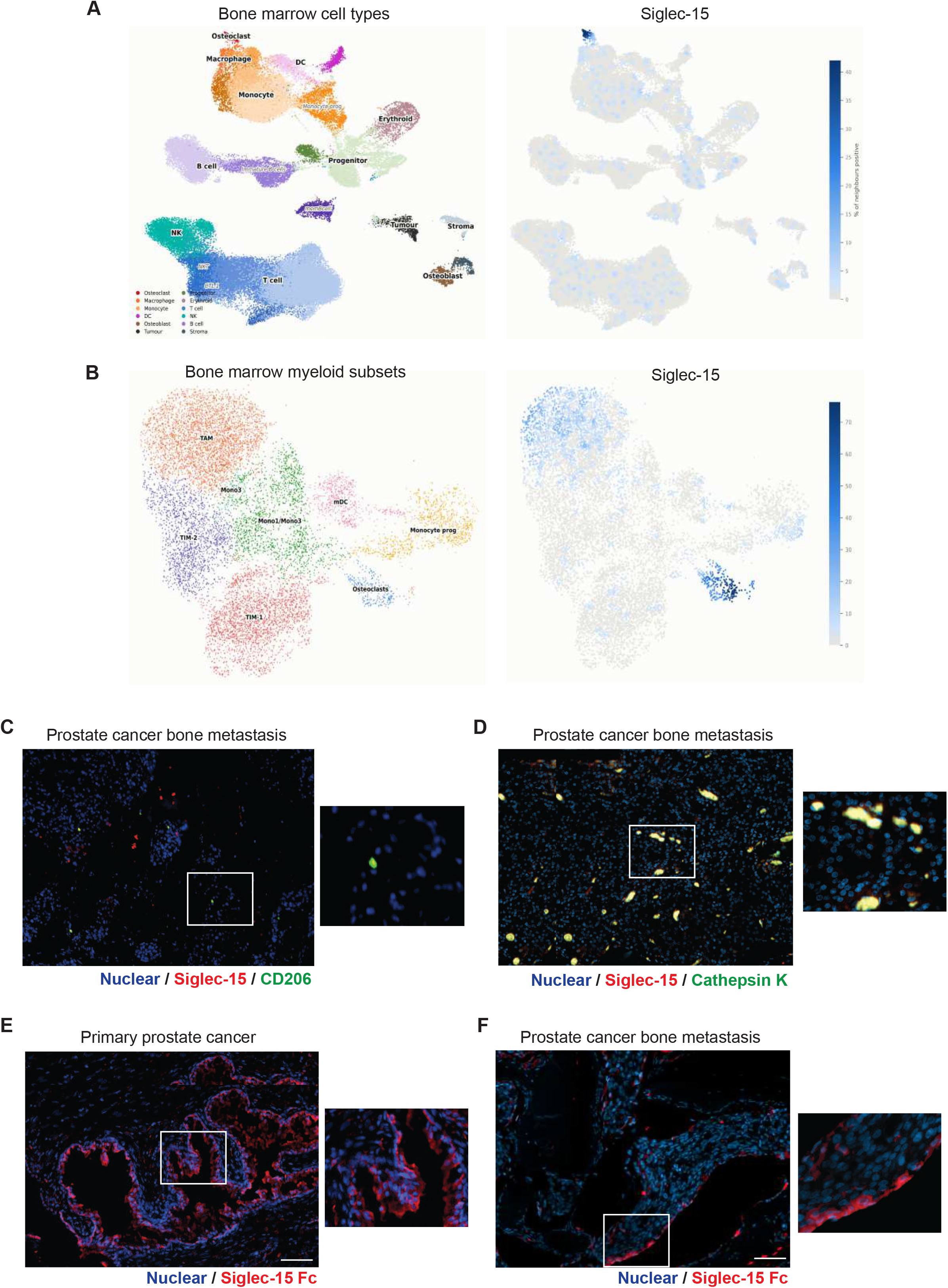
Siglec-15 is expressed by prostate cancer cells, M2 macrophages and osteoclasts in the prostate cancer tumour immune microenvironment. **(A,B)** UMAP map of previously published single cell RNA-sequencing datasets from bone metastatic prostate tumours (37, 38) showing Siglec-15 expression levels across different cell types. Interrogation of the intra-tumour myeloid sub-compartments within these combined datasets shows SIGLEC15 transcripts are expressed by osteoclasts and tumour-associated macrophage (TAM) subsets. **(C,D)** Dual immunofluorescence analysis of Siglec-15 and immune cell markers in prostate-derived tumours growing in the bone. Siglec-15 is co-expressed with the M2 macrophage marker CD206 and cathepsin K (an osteoclast marker). **(E,F)** Siglec-15 sialoglycan ligands are expressed in primary prostate tumours and in bone metastatic prostate tumours (monitored using Siglec-15 Fc immunofluorescence). Scale bar is 100 µm.

For the transcriptomic analysis in Figure 4, we analysed the TGCA-PRAD dataset (41) using methods described by Gyorffy et al (42). We scored for the expression of a combination of genes, determined using the Siglec-15 dependent significant differentials from our proteomic data (flow and ELISA assays) which showed significant differences when comparing treatments (control or anti-Siglec-15 antibodies). This gave us a Siglec-15+, Siglec-15 dependent macrophage (SSDM) transcriptomic signature of *SIGLEC15*, *MRC1*, *TGFB1*, *CD274* and *CD86*. We then normalised the expression of CD8A for each case using *PTPRC* (*CD45*) and split normalised CD8A expression into two groups: those cases with high or low SSDM scores (median split). Finally, to assess the impact of these cells on survival, we split the SSDM scores into top and bottom thirds and plotted the relevant prognostic data for each case.

### Statistical analyses

Statistical analyses were performed using GraphPad Prism (Version 11.0.2; GraphPad Software). Data were assessed for normality prior to analysis, and homogeneity of variance was evaluated using either an F-test or Levene’s test, as appropriate. Statistical tests were selected based on the data distribution and experimental design, as specified in the corresponding figure legends and main text. Data are presented as the mean ± standard error of the mean (SEM) from three independent biological replicates. Statistical significance was defined as p < 0.05, **p < 0.01, *** p < 0.001, and **** p < 0.0001.

## RESULTS

### 1. Siglec-15 is expressed by prostate tumour cells, M2 macrophages, and osteoclasts within the prostate tumour microenvironment

Given the emerging role of Siglec-15 as a critical glyco-immune checkpoint, we first sought to map its spatial and cellular distribution within the prostate tumour microenvironment. Having previously characterized *SIGLEC15* expression at single-cell resolution across published single-cell RNA-sequencing datasets (37, 43), we established that *SIGLEC15* transcripts reside predominantly within myeloid cells, macrophages, and osteoclasts (20). To gain deeper resolution into the bone metastatic niche, we expanded our initial analysis of the index bone metastasis dataset (37), by integrating a second independent bone metastasis cohort (38). This integrated analysis confirmed that *SIGLEC15* is highly enriched within osteoclasts, consistent with its established role in bone remodelling (**Figure 1A**). Furthermore, high resolution interrogation of the intra-tumour myeloid sub-compartments within these combined datasets revealed that *SIGLEC15* transcripts are also expressed by tumour-associated macrophage (TAM) subsets (**Figure 1B**).

To validate these transcriptomic findings at the protein level, we performed dual immunofluorescence analysis of whole slide sections from 5 primary prostate tumours and 5 bone metastatic lesions. Siglec-15 protein expression co-localised with the malignant epithelial marker α-methylacyl-CoA racemase (AMACR) in both primary and metastatic tumours (Supplementary Figure 1). Furthermore, Siglec-15 was detected on CD206⁺ M2-polarised macrophages in primary tumours (Supplementary Figure 1), as well as on both multinucleated osteoclasts and CD206⁺ TAMs within the bone metastatic tissue (**Figure 1C,D**). Finally, to assess the presence of Siglec-15 ligands within the tumour microenvironment, we performed immunofluorescence staining using a Siglec-15 Fc protein. This demonstrated widespread presence of Siglec-15 binding sialoglycans in both primary and bone metastatic lesions (**Figure 1E,F**). Taken together, these findings establish that Siglec-15 and its glycan ligands are co-expressed across malignant, immune, and bone-resorbing cell types in prostate cancer, supporting a potential role for the Siglec-15 sialoglycan axis in tumour-associated immune suppression and osteoclast mediated bone metastasis.

### 2. Siglec-15 is upregulated in prostate tumour epithelial cells

Having identified Siglec-15 expression in prostate tumour tissues, we next investigated whether its expression is altered during prostate cancer progression. Although Siglec receptors are classically associated with immune cells, accumulating evidence indicates that several family members, including Siglec-15, can also be expressed by tumour cells (26, 44–46). To evaluate Siglec-15 expression in tumour epithelial cells, we utilised a well validated immunohistochemical assay that was previously developed as a potential companion diagnostic for clinical trials (33). Using Siglec-15 immunohistochemistry, we analysed three independent prostate cancer tissue microarray (TMA) cohorts comprising normal prostate tissue and primary prostate cancers. Analysis of a TMA containing tissue samples from 40 prostate cancer patients (TMA cohort 1) showed that Siglec-15 is expressed at significantly higher levels in prostate cancer tissue compared to normal prostate tissue (Supplementary Figure 2). Further analysis of a 96 case TMA containing 17 normal prostate tissue samples and 79 samples of prostate tumour tissue (TMA cohort 2) also showed that Siglec-15 is significantly upregulated in prostate tumours (unpaired t test, p <0.0001) (**Figure 2A**).

**Figure 2.**
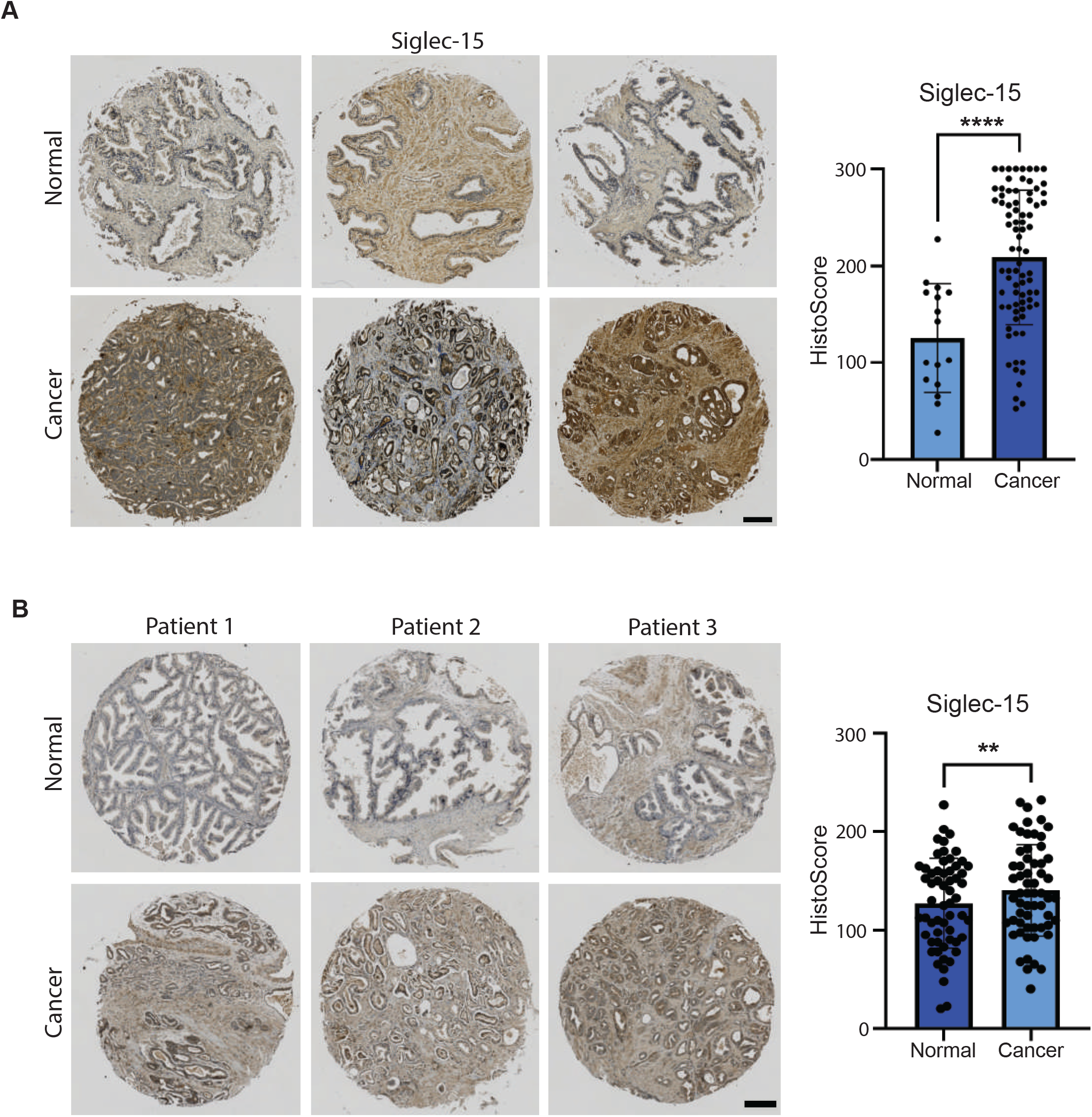
Siglec-15 is upregulated in prostate tumour epithelial cells. (**A**) Immunohistochemistry analysis of Siglec-15 in a previously published TMA (19) comprising 96 prostate tissue samples shows Siglec-15 is detected at significantly higher levels in prostate tumour tissue relative to normal prostate tissue (unpaired t test, p < 0.0001). (**B**) Immunohistochemistry analysis of Siglec-15 in a TMA comprising tissue from 60 cases of prostate cancer and matched normal tissues (TMA cohort 3). Siglec-15 is detected at significantly higher levels in in prostate cancer cells relative to matched normal prostate tissue (paired t test, p=0.0173, only epithelial cells were scored). Scale bar is 100 µm.

Staining of a TMA comprising matched normal and prostate cancer tissue from 60 patients (TMA cohort 3) showed that Siglec-15 is found at significantly higher levels in prostate cancer epithelial cells relative to matched normal prostate tissue from the same patient (paired t test, p = 0.0173) (**Figure 2B**). Collectively, these independent patient cohorts demonstrate that Siglec-15 is consistently upregulated in prostate tumour epithelial cells, identifying tumour cell expression of Siglec-15 as a feature of primary prostate cancer.

### 3. Siglec-15 is enriched in bone metastatic prostate cancer and maintained across advanced CRPC subtypes

The majority of patients with advanced prostate cancer receiving hormone therapy eventually progress to CRPC, a disease state frequently accompanied by metastatic progression. To investigate whether Siglec-15 expression is associated with metastatic CRPC, we analysed a previously described TMA comprising matched bone and visceral (liver) metastases collected at rapid autopsy from 100 patients with lethal CRPC (20) (TMA cohort 4). Immunohistochemical analysis demonstrated that Siglec-15 expression was significantly higher in bone metastases than in matched visceral metastases from the same patients (paired t-test, p<0.0002), suggesting enrichment of Siglec-15 within the prostate cancer bone metastatic niche (**Figure 3A**). To determine whether Siglec-15 expression is maintained across clinically relevant models of advanced CRPC, we utilised samples from the LuCaP patient derived xenograft (PDX) series (31, 32). Using immunohistochemistry, we evaluated Siglec-15 expression in a TMA comprising 39 LuCaP PDX models established in intact and castrated immunodeficient mice. This platform comprehensively captures the molecular and phenotypic heterogeneity of advanced CRPC, including AR positive adenocarcinoma, neuroendocrine prostate cancer (NEPC) and double negative prostate cancer (DNPC) phenotypes. Siglec-15 was detected across both AR-positive, NEPC, and DNPC models, demonstrating that expression is maintained across distinct molecular subtypes of advanced prostate cancer (**Figure 3B**). To complement these protein level findings, we leveraged data from published NEPC gene datasets (36). Integrated analysis of single-cell and bulk transcriptomic datasets, revealed that NEPC-derived tumour cells exhibit high intrinsic expression of *SIGLEC15*, suggesting that this glyco-immune checkpoint is a feature of highly aggressive, transdifferentiated neuroendocrine lineages (Supplementary Figure 3). Furthermore, immunohistochemistry profiling showed Siglec-15 expression is similar between tumours grown in intact and castrated hosts, suggesting that its expression is not substantially altered by androgen status (**Figure 3C**). Together, these findings demonstrate that Siglec-15 is enriched in prostate cancer bone metastases and is expressed across diverse, treatment resistant molecular landscapes of advanced disease.

**Figure 3.**
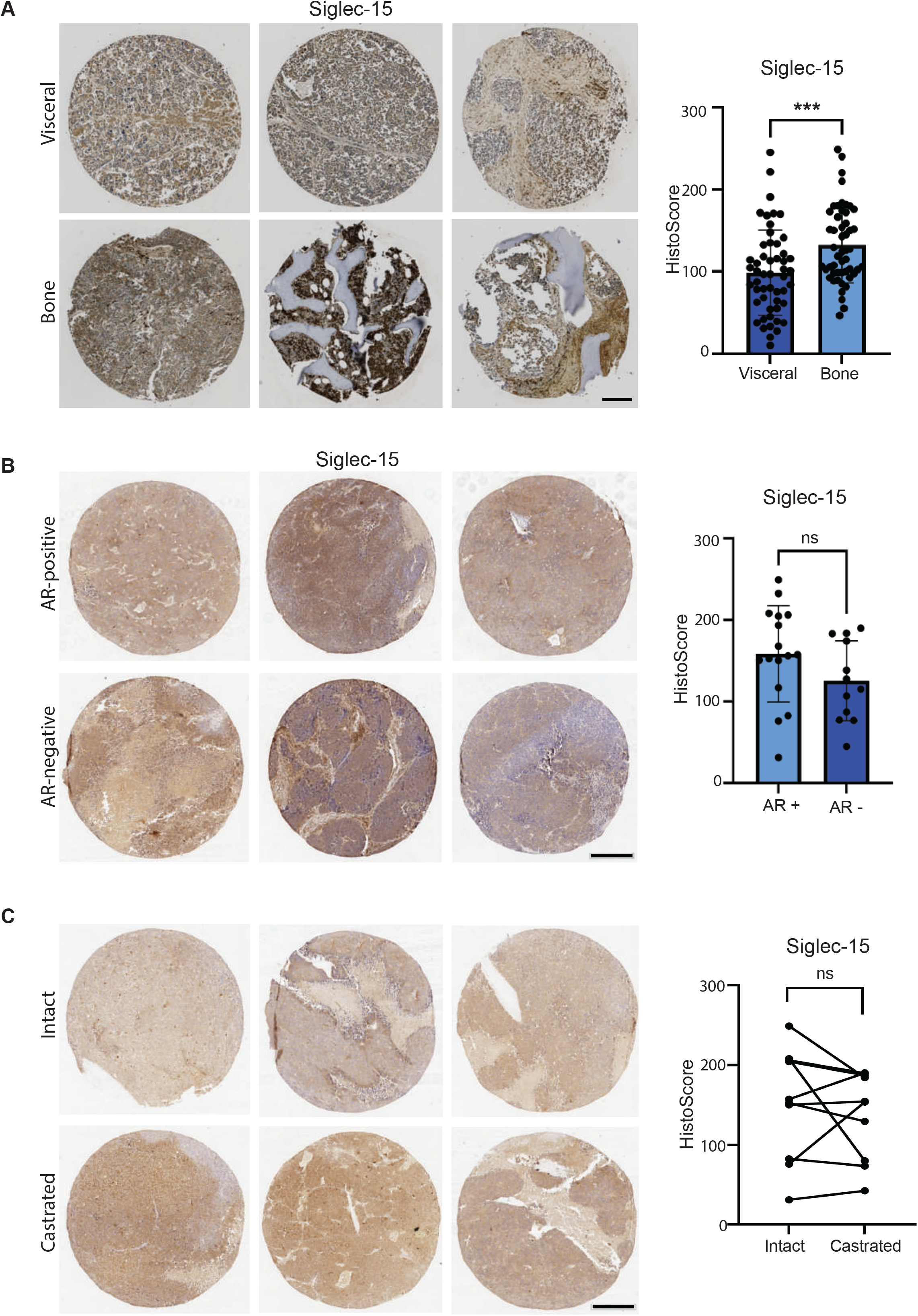
Expression of Siglec-15 in CRPC prostate cancer. (**A**) Immunohistochemistry analysis of Siglec-15 in a TMA containing matched visceral and bone CRPC metastatic tumours from 100 patients obtained via rapid autopsy (64). Siglec-15 Histoscores were significantly higher in prostate-derived tumours growing in bone compared to matched visceral tumour tissue from the same patient (n=100, paired t test, p<0.0002, only epithelial cells were scored). Scale bar is 200 µm. (**B**) Analysis of Siglec-15 in LuCaP patient-derived xenografts (PDX) tissues. Immunohistochemistry was used to monitor Siglec-15 in a TMA comprising 39 samples from the LuCaP PDX series (31, 32). Siglec-15 Histoscores showed no significant change in Siglec-15 levels in AR-null (neuroendocrine) prostate cancer models compared to AR-positive prostate cancer models grown in intact mice (n=28, unpaired t test, p=0.1188). (**C**) Comparison of Siglec-15 levels in PDX tissues established from the same patient grown in intact or castrated immunocompetent mice. There was no significant change in Siglec-15 levels when comparing intact or castrated conditions (paired t rest, p=0.4936). Representative images for PDX models are shown. Each PDX sample was analysed in triplicate within 3 mice. Scale bar is 200 µm.

### 4. Siglec-15 regulates monocyte differentiation and promotes prostate tumour growth

Having established that Siglec-15 is expressed by tumour cells and immunosuppressive myeloid populations within the prostate tumour immune microenvironment, we first investigated the cell autonomous role of Siglec-15 activation in early myeloid development. We first confirmed that Siglec-15 is expressed by primary human monocytes and macrophages by flow cytometry (**Figure 4A**). To determine whether direct engagement of this receptor actively drives monocyte fate, primary human peripheral blood monocytes were cultured in the presence of humanised anti-human Siglec-15 (Clone NC318) and goat anti-human Fc to induce functional receptor crosslinking. Siglec-15 crosslinking alone was sufficient to drive monocyte differentiation, suggesting that ligand-mediated crosslinking of Siglec-15 acts as an inductive switch that can actively drive monocyte differentiation (**Figure 4B**). Next, to closely mirror the complex secretory profile of the localised prostate tumour microenvironment, we cultured primary human peripheral monocytes in concentrated conditioned media (CM) collected from CWR22Rv1 prostate cancer cells to drive differentiation toward tumour-associated macrophages. Treatment with the anti-Siglec-15 antibody (Clone 5G12, a parent of the humanised clinical-stage anti-human Siglec-15 antibody NC318) significantly decreased the number of live macrophages within these cultures (**Figure 4C**). This indicates that the Siglec-15 axis is required to support myeloid cell survival or expansion in response to the tumour secretome. We next investigated whether these immune-modulatory effects translated into anti-tumour activity *in vivo* using the immunocompetent TRAMP-C2 syngeneic mouse model. Systemic treatment with the anti-Siglec-15 antibody (Clone 5G12, NextCure) significantly suppressed subcutaneous tumour growth over 75 days (p <0.0001) (**Figure 4D**). Importantly, depletion of CD8⁺ T cells, in line with the reported effects of Siglec-15+ macrophages (26), abolished the anti-tumour effects of Siglec-15 blockade, demonstrating that the therapeutic efficacy of Siglec-15 inhibition is dependent upon cytotoxic T cell responses. Collectively, these findings demonstrate that Siglec-15 receptor crosslinking acts as a driver of monocyte differentiation, a mechanism that may be exploited by factors secreted by prostate cancer cells. Furthermore, therapeutic blockade of the Siglec-15 axis remodels this myeloid niche to restrict prostate tumour growth via a CD8⁺ T-cell-dependent mechanism.

**Figure 4.**
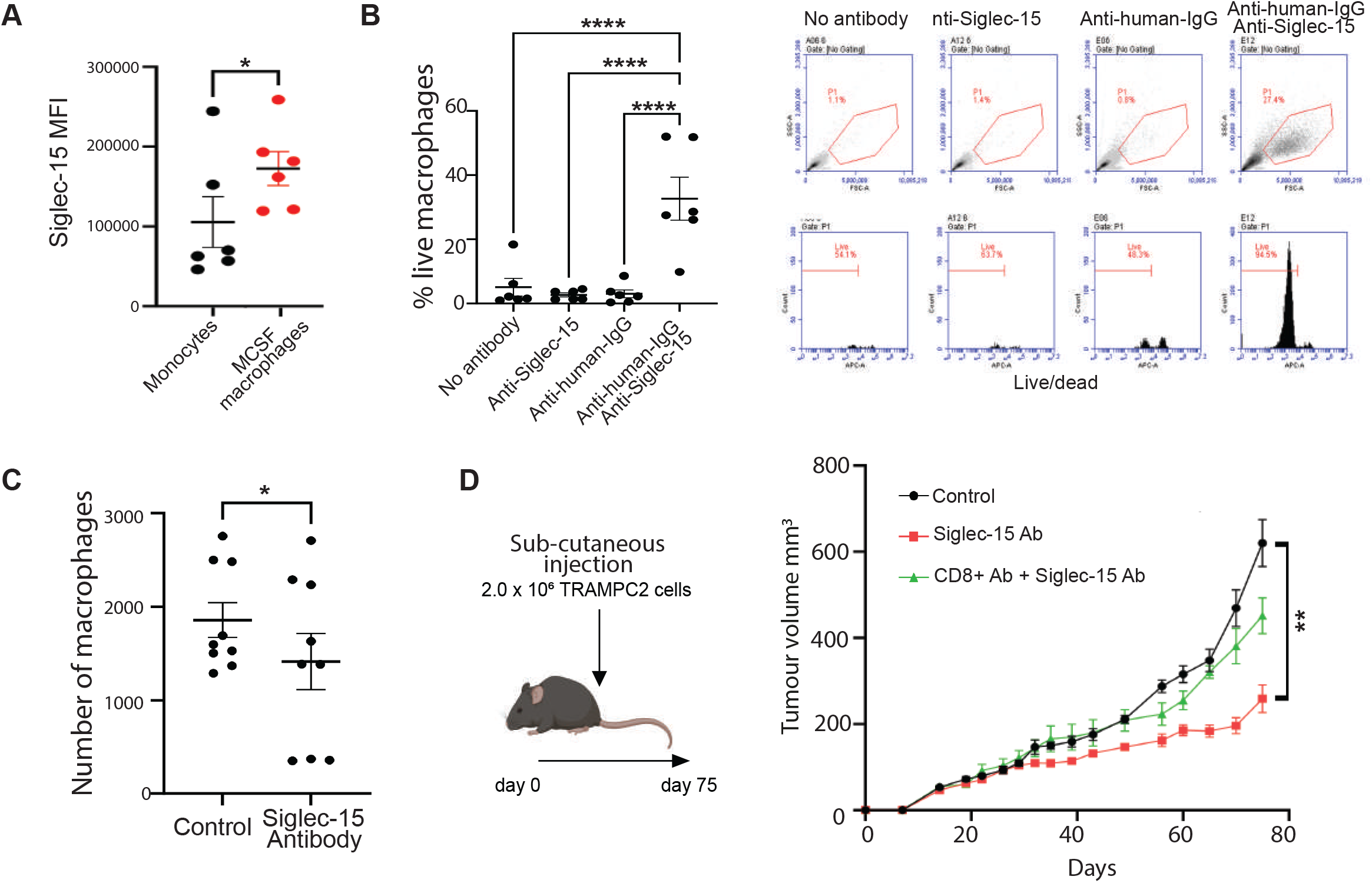
Targeting Siglec-15 suppresses prostate tumour growth through CD8⁺ T-cell-dependent immunity. (**A**) Siglec-15 is expressed by healthy primary monocytes and M-CSF monocyte-derived macrophages (detected using NC318, NextCure). N=6 (Biological). (**B**) Crosslinking siglec-15 (NC318 plus anti human IgG) drives monocyte differentiation. N=6 (Biological). Example plots shown. (**C**) Concentrated supernatant from CWR22Rv1 cells drives monocyte differentiation. N=3 (Biological) N=3 (Technical) (**D**) Concentrated supernatant from CWR22Rv1 cells drives monocyte differentiation in a partially siglec-15 dependent manner. N=3 (Biological) N=3 (Technical). (**D**) Schematic of study design for TRAMP-C2 allograft model. 2×10^6^ TRAMP-C2 cells were injected sub-cutaneously into the left flank of C57BL/6 mice. Tumour growth was monitored using callipers over 75 days. Tumour growth curves for TRAMP-C2 allografts in control, anti-CD8 and anti-Siglec-15, and anti-Siglec-15 (5G12, NextCure) treated groups (6 mice per group). Tumour growth was significantly reduced in mice treated with anti-Siglec-15 antibodies compared to the control group, but not in mice treated with both anti-CD8 and anti-Siglec-15 antibodies (Two-way ANOVA, p <0.0001).

### 5. Siglec-15 coordinates M-CSF-driven macrophage programming with CD8⁺ T-cell suppression and osteoclastogenesis in prostate cancer bone metastasis

Given that advanced prostate cancer has a high propensity to metastasise to bone, we next investigated how cytokines present in the bone metastatic microenvironment interface with the Siglec-15 pathway. M-CSF is constitutively expressed in the bone marrow, highly elevated during bone colonisation and is a known driver of the ‘vicious cycle’ (47, 48). We therefore evaluated the impact of M-CSF on primary monocyte differentiation, macrophage phenotype and CD8 T^+^ cell suppression. Primary human peripheral blood monocytes were cultured in the presence of recombinant M-CSF to induce macrophage differentiation. While monocyte exposure to M-CSF drove differentiation into a well characterised immunosuppressive cell phenotype, the introduction of the anti-Siglec-15 blocking antibody had little effect on the % of live macrophages (**Figure 5A**), instead altering M-CSF-driven macrophage polarisation, leading to a significant reduction in surface CD206, PD-L1 and CD86 expression (**Figure 5B**) alongside reduced secretion of TGF-β1 and an increase in proliferating PBMCs (**Figure 5C** and Supplementary Figure 4). Functionally, supernatant from both cell line and MCSF monocyte derived macrophages suppressed CD8⁺ T-cell proliferation and IFN-γ production, effects that were partially restored following Siglec-15 blockade during macrophage differentiation (**Figure 5D,E**).

**Figure 5.**
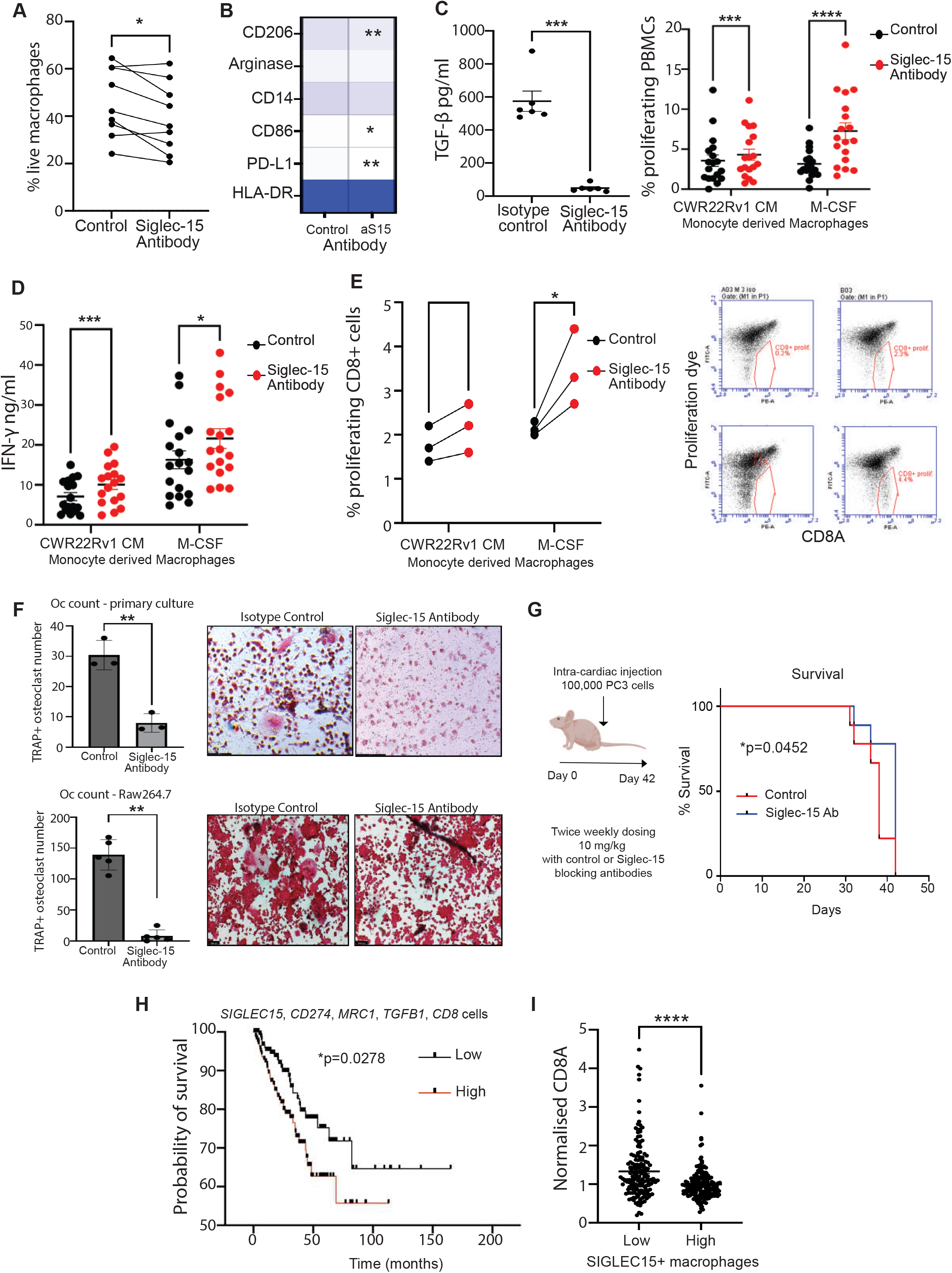
Targeting Siglec-15 inhibits M-CSF pathways and osteoclastogenesis, and prolongs survival in a mouse model of bone metastasis. (**A**) M-CSF drives the differentiation of monocytes into macrophages in a partially Siglec-15 dependent manner. N=3 (Biological), N=3 (Technical). (**B**) M-CSF drives the differentiation of monocytes into immunosuppressive macrophages (CD206+, PD-L1+, CD86+) in a partially Siglec-15 dependent manner. N=6 (Biological). (**C**) The concentration of total TGF-β present in the supernatant from cells from B N=3 (Biological) N=2 (Technical). (**D,E**) Supernatant from cells from Figure 4C, and Figure 5B inhibited the CD3/CD28 mediated proliferation and IFN-y production of PBMCs. Example plots shown. (**F**) Treatment with anti-Siglec-15 antibody significantly inhibits osteoclast differentiation, as determined by tartrate-resistant acid phosphatase (TRAP) staining (tested using primary murine osteoclast precursors and mouse RAW264.7 cells). Scale bar is 100 µm. (**G**) Schematic of study design for PC3 intra-cardiac bone metastasis model. Balb/c nude mice were injected with 100,000 PC3 cells. Twice weekly treatment with 10mg/kg control (NP149, NextCure) or anti-Siglec-15 antibodies (NP159, Fc silenced, NextCure) via intraperitoneal (IP) injection significantly prolonged survival compared with control treated mice (n=20, Log-rank test, p = 0.0452). (**H**) Analysis of the TCGA PRAD cohort (41) identifies a Siglec-15 dependent macrophage signature. Prostate cancer tumours with a high number of *SIGLEC15*, *CD274*, *CD86*, *MRC1*, *TGF*β macrophages have a significantly worse prognosis (p=0.0278). (**I**) Analysis of the TCGA PRAD cohort (41) shows macrophages with high SIGLEC15 levels negatively correlate with CD8A number and function.

As Siglec-15 is highly expressed by osteoclasts within prostate cancer bone metastases, we investigated whether therapeutic blockade of Siglec-15 could inhibit osteoclast differentiation and tumour progression in bone. We first confirmed that Siglec-15 is expressed by primary human osteoclasts by demonstrating specific binding of anti-Siglec-15 antibodies, confirming expression of Siglec-15 within the osteoclast compartment and supporting its potential role in bone metastatic disease (Supplementary Figure 5). To determine whether Siglec-15 contributes to tumour-induced osteoclastogenesis, primary mouse osteoclast pre-cursors and mouse RAW264.7 cells were differentiated into osteoclasts in the presence or absence of anti-Siglec-15 antibody. Treatment with anti-Siglec-15 antibody (Clone NC318) significantly reduced the formation of TRAP-positive osteoclasts, suggesting that therapeutic blockade of Siglec-15 can inhibit tumour-induced osteoclast differentiation (**Figure 5F**). Next, to investigate whether these findings translate into therapeutic benefit, we utilised a mouse model of prostate cancer bone metastasis established by intra-cardiac injection of PC3 prostate cancer cells. Treatment with anti-Siglec-15 antibody significantly prolonged survival compared with control treated mice, suggesting that therapeutic inhibition of Siglec-15 suppresses metastatic disease progression (p=0.0452) (**Figure 5G**). To validate these translational findings in human cohorts, we developed a gene expression signature representing the Siglec-15-dependent immunosuppressive macrophage phenotype identified in our *in vitro* assays (*SIGLEC15*, *CD274*, *CD86*, *MRC1*, and *TGFB1*). Prognostic evaluation revealed that prostate cancer patients with a high enrichment score for this myeloid signature experienced significantly worse survival outcomes compared to those with low expression (p = 0.0278, **Figure 5H**). Finally, to determine the environmental impact of these specific macrophages on local T-cell infiltration *in situ*, we correlated our macrophage signature score against tumour CD8A expression. Consistent with our co-culture data, we observed a significant negative correlation between the Siglec-15-dependent macrophage signature and CD8A levels (**Figure 5J**), confirming that this myeloid axis correlates with an excluded or suppressed T cell microenvironment in clinical disease. Together, these findings position Siglec-15 as a central regulator of the metastatic myeloid niche, linking M-CSF-driven macrophage programming with CD8⁺ T-cell suppression and osteoclastogenesis, and highlight Siglec-15 blockade as a potential therapeutic strategy in metastatic prostate cancer.

## Discussion

Despite substantial advances in the treatment of prostate cancer, metastatic CRPC remains a major clinical challenge, with limited therapeutic options and poor long-term outcomes (49). Although immune checkpoint blockade has transformed the treatment of several cancer types, responses in prostate cancer have been disappointing (50) highlighting the need to identify alternative mechanisms of tumour immune evasion (51). In this study, we identify Siglec-15 as a previously unrecognised regulator of prostate cancer progression that links tumour-associated immune suppression and bone metastatic disease. We demonstrate that Siglec-15 is expressed by ‘M2’ immunosuppressive macrophages, osteoclasts, and prostate cancer epithelial cells within the prostate tumour microenvironment, is enriched in bone metastatic CRPC, and promotes both macrophage-mediated immune suppression and osteoclast activity. Importantly, therapeutic blockage of Siglec-15 suppresses tumour growth and prolongs survival in pre-clinical models, supporting Siglec-15 as a promising therapeutic target for advanced prostate cancer.

Aberrant tumour sialylation is increasingly recognised as a hallmark of cancer progression, with cancer-associated sialoglycans engaging inhibitory Siglec receptors to establish an immunosuppressive tumour microenvironment (8). This glyco-immune checkpoint represents an important mechanism of immune escape that operates independently of established pathways such as PD-1/PD-L1 (52, 53). Our findings build upon a growing body of work demonstrating that prostate cancer undergoes extensive glycan remodelling during disease progression (15, 20, 54). We recently showed that prostate tumours upregulate Siglec-engaging immunosuppressive sialoglycans that promote tumour progression and bone metastasis, and that disruption of these glycan-mediated interactions remodels the tumour immune microenvironment by reducing immunosuppressive macrophages while enhancing natural killer (NK) cell cytotoxicity (20). More recently, we identified the cancer-associated sialyl-Tn (sTn) antigen as a clinically relevant component of this altered glycome, demonstrating that sTn expression is associated with adverse clinical outcomes and is retained in therapy resistant metastatic disease, including bone metastases (19). This study builds on these findings by identifying Siglec-15 as a key functional effector of this glyco-immune network. We have demonstrated that direct Siglec-15 receptor crosslinking operates as an inductive switch that actively drives monocyte differentiation. Crucially, we found that M-CSF, a growth factor heavily enriched within the prostate and bone marrow niches (55), can actively exploit this axis to program monocytes into an immunosuppressive state. This suggests that instead of simply blocking T cell activation on the surface, Siglec-15 may actively signal to drive the development and differentiation of monocytes into immunosuppressive macrophage phenotypes (**Figure 6**).

**Figure 6.**
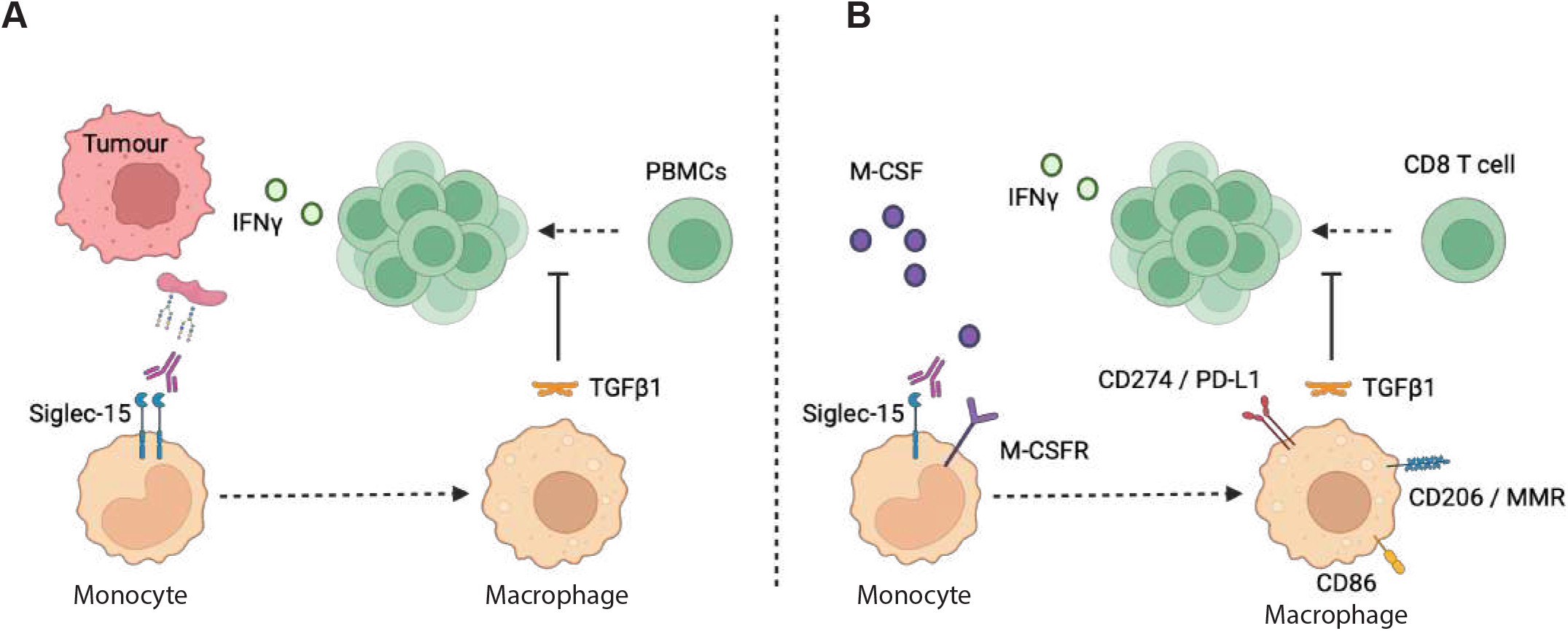
Schematic of putative Siglec-15 dependent mechanism of monocyte-derived macrophage generation and associated immunological impacts. (**A**) Tumours (red) produce secreted hypersialylated structures which are able to cross-link Siglec-15 on monocytes (beige). This Siglec-15 dependent crosslinking results in activation and differentiation into TGFb1highmacrophages (beige) which inhibit PBMC proliferation (green) and IFNy release. (**B**) MCSF (from the local environment) drives the generation of monocyte-derived macrophages in a partially Siglec-15 dependent manner, most likely through a ‘yet to be determined’ cis-interaction. This Siglec-15 dependent signalling results in activation and differentiation into TGFb1high, PD-L1high, CD206high, CD86high macrophages which inhibit CD8 T cell proliferation and IFNy release. As such, the inhibition of these processes through the use of therapeutic Siglec-15 blockade may result in improved prognosis.

Although early studies proposed the cancer-associated sTn antigen as a ligand for Siglec-15, more recent investigations have reported conflicting findings regarding the glycan-binding specificity of Siglec-15 (56–60), and its physiologically relevant endogenous ligands remain to be fully defined. Importantly, our demonstration that Siglec-15 is expressed across multiple cell types, including prostate cancer epithelial cells, ‘M2’ macrophages and osteoclasts, expands the role of this pathway beyond a conventional immune checkpoint. Rather, Siglec-15 may function as a multicellular regulator of the metastatic niche, coordinating interactions between tumour cells, immune populations and bone-resident cells to promote immune suppression and osteoclast-mediated bone remodelling. Although the precise tumour-intrinsic functions of Siglec-15 remain to be determined, epithelial tumour cell expression suggests that in addition to regulating immune cell function, Siglec-15 may contribute directly to tumour-microenvironment communication. Together with our previous studies (11, 19, 20), the data presented here support a model in which coordinated alterations in tumour sialylation and Siglec signalling promote prostate cancer progression and identify the Siglec-15 sialoglycan axis as a promising therapeutic vulnerability in advanced disease. Interestingly, while antibody blockade did not alter the expression of the phenotypic surface marker CD206 within tumour conditioned cultures, it significantly decreased the absolute number of live macrophages. This suggests that the prostate tumour secretome may utilise Siglec-15 to polarise macrophages, and to actively sustain their survival and expansion, through a putative sialylated crosslinking agent. Furthermore, the remaining macrophage pool exhibited a reduction in the secretion of the immunosuppressive and pro-fibrotic cytokine TGF-β. This may be particularly relevant in prostate cancer, which is generally regarded as an immunologically cold tumour that exhibits a limited response to conventional immune checkpoint inhibitors (50, 51).

In addition to its immunological functions, accumulating evidence has highlighted an important role for Siglec-15 in osteoclast differentiation and bone remodelling (22, 23, 61). In breast cancer, the Siglec-15 sialoglycan axis has been proposed as a central regulator linking immune suppression with osteoclast mediated bone destruction during metastatic progression (25). Our findings extend this concept to prostate cancer, which has a marked propensity for skeletal dissemination. We show that Siglec-15 is expressed by osteoclasts within prostate cancer bone metastases and that antibody blockade suppresses osteoclast differentiation while significantly prolonging survival in mice with prostate cancer bone metastasis. These findings suggest that Siglec-15 inhibition may provide dual therapeutic benefit by simultaneously restoring anti-tumour immunity and limiting osteoclast driven remodelling of the metastatic bone microenvironment. Such dual targeting may represent a particular advantage over therapies that act exclusively on either tumour immunity or bone turnover.

Our finding that Siglec-15 is expressed in both AR positive adenocarcinoma models and AR negative NEPC models suggests that its expression is not restricted to a single molecular subtype of advanced disease. Given the increasing clinical importance of treatment emergent CRPC phenotypes which are associated with poor prognosis and limited therapeutic options (62, 63), these findings raise the possibility that Siglec-15 directed therapies may have broad applicability across multiple advanced prostate cancer phenotypes. Future studies should determine whether Siglec-15 blockade can be effectively combined with existing therapies, including AR pathway inhibitors, immune checkpoint inhibitors and bone directed treatments.

Several important questions remain. Although our findings establish a critical role for Siglec-15 in directly regulating macrophage differentiation and indirectly regulating CD8⁺ T cell-dependent tumour immunity, the downstream molecular pathways linking Siglec-15 and M-CSF signalling to adaptive immune activation remain incompletely understood. Given the relative lack of impact of a) M-CSF on Siglec-15 crosslinked macrophage numbers, and b) anti-Siglec-15 on M-CSF macrophage numbers, there may be evidence of shared differentiating signalling, most likely through DAP12. Furthermore, although we demonstrate inhibition of osteoclast differentiation and improved survival in a mouse models of prostate cancer bone metastasis, future studies incorporating spatial multi-omics and glycomic profiling will be important for defining how prostate tumour cells, immune cells and bone resident cells cooperate to establish the Siglec-15-dependent metastatic niche. Finally, the endogenous glycoprotein carriers presenting the sialoglycan ligands that regulate Siglec-15 signaling in prostate cancer have not yet been identified. Defining these ligand-receptor interactions will not only improve our understanding of prostate cancer glycoimmunology but may also reveal additional therapeutic opportunities.

In conclusion, our findings establish Siglec-15 as a dual regulator of tumour-associated immune suppression and osteoclast bone pathology in prostate cancer. Together with our previous studies demonstrating remodelling of the prostate cancer sialoglycome (12, 14, 15, 20), these data support a model in which coordinated alterations in tumour glycosylation and Siglec signalling establish a glyco-immune axis that promotes immune evasion and metastatic progression. Therapeutic targeting of the M-CSF sialoglycan-Siglec-15 axis therefore represents a promising strategy to simultaneously enhance anti-tumour immunity and disrupt the bone metastatic microenvironment in patients with advanced prostate cancer.

## Supplementary Figures

**Supplementary Figure 1. Siglec-15 colocalises with AMACR in primary prostate tumours and prostate cancer bone metastasis and is expressed by M2 macrophages in primary prostate tumours.** (**A,B**) Dual immunofluorescence analysis shows Siglec-15 is co-expressed with AMACR and CD206 in primary prostate cancer (confirming Siglec-15 expression within cancerous tissues and by immunosuppressive macrophages in prostate tumours). Scale bar is 100 µm. (**C**) Dual immunofluorescence analysis shows Siglec-15 is co-expressed with AMACR in prostate cancer bone metastasis (confirming Siglec-15 expression within cancerous tissues). Scale bar is 50 µm.

**Supplementary Figure 2. Siglec-15 is upregulated in prostate tumour epithelial cells.** (**A**) Immunohistochemistry analysis of Siglec-15 in a previously published TMA (20) comprising 51 prostate tissue samples (TMA cohort 1) shows Siglec-15 is detected at significantly higher levels in prostate tumour tissue relative to normal prostate tissue (unpaired t test, p=0.0093). Scale bar is 100 µm.

**Supplementary Figure 3.** Analysis of integrated datasets from primary prostate tumours (36) shows *SIGLEC15* is highly expressed in NEPC tumour cells (labelled NE).

**Supplementary Figure 4.** Supernatant from cells from Figure 4C, and Figure 5B inhibited CD3/CD28 mediated CD8+ cell proliferation in a Siglec-15 dependent manner. Example plots shown.

**Supplementary Figure 5.** Siglec-15 is expressed by osteoclasts differentiated from human monocytes (detected using NC318).

## ACKNOWLEDGEMENTS

This work was funded by the Prostate Cancer Research and Breast Cancer Now bone metastasis collaboration fund (grant reference BMCF01), the JGW Patterson Foundation, Prostate Cancer Research (grant reference 6974), Prostate Cancer UK [RIA21-ST2-006] and the Medical Research Council [MR/R015902/1]. This work is also supported by the Pacific Northwest Prostate Cancer SPORE (P50CA97186), the PO1 NIH grant (PO1 CA163227), the Prostate Cancer Foundation, and the Institute for Prostate Cancer Research (IPCR).

## Data availability

The authors confirm that the data supporting the findings of this study are available within the article and its supplementary materials.

