## Supplementary Figures for "Siglec-15 is a glyco-immune checkpoint in prostate cancer regulating immune evasion and metastasis"

Supplementary Figure 1

A

Primary prostate cancer

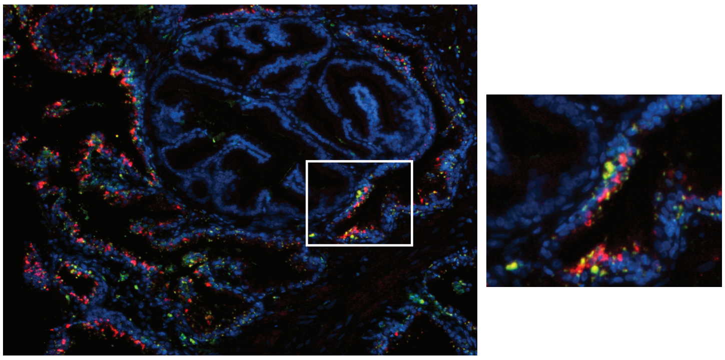

Nuclear / Siglec-15 / AMACR

B

Primary prostate cancer

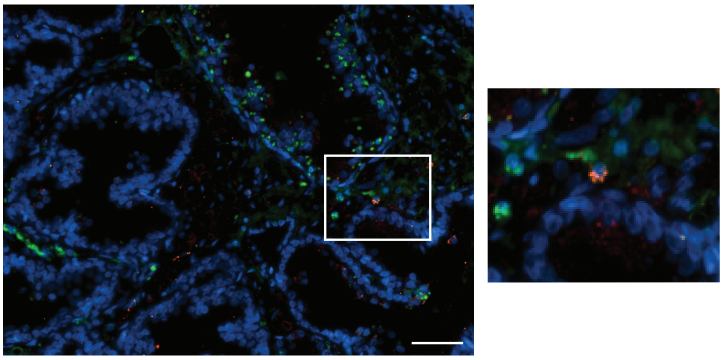

Nuclear / Siglec-15 / CD206

C

Prostate cancer bone metastasis

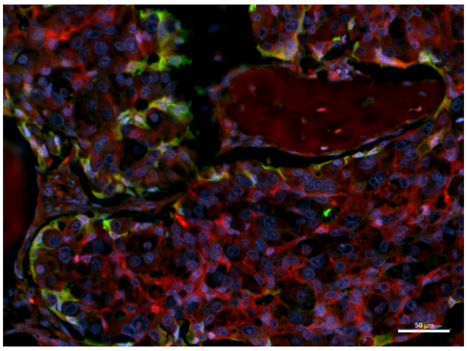

Nuclear / AMACR / Siglec-15

Supplementary Figure 2

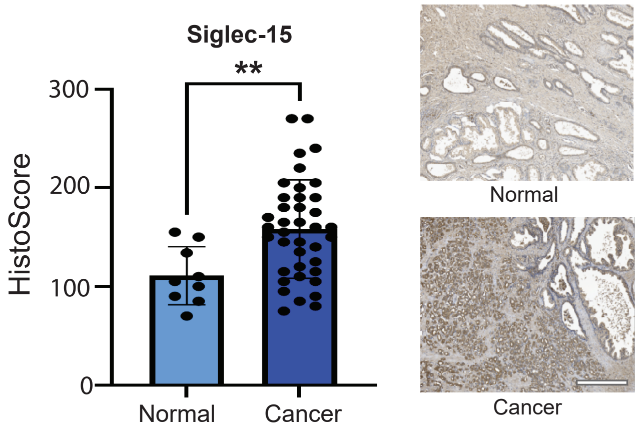

Supplementary Figure 3

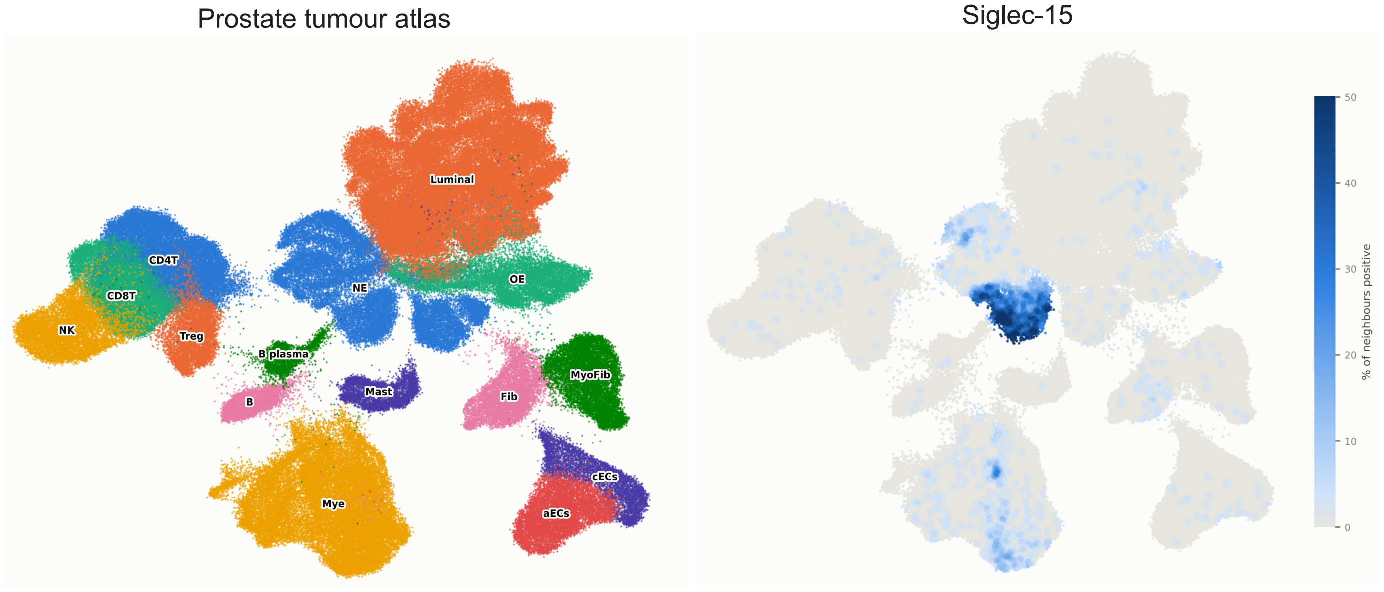

Supplementary Figure 4

A

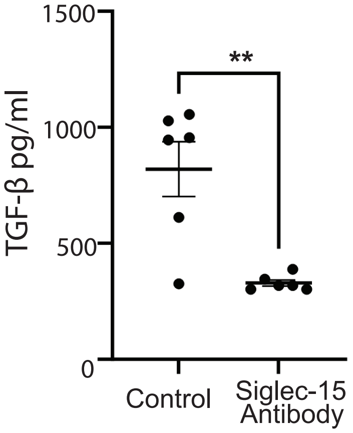

Supplementary Figure 5

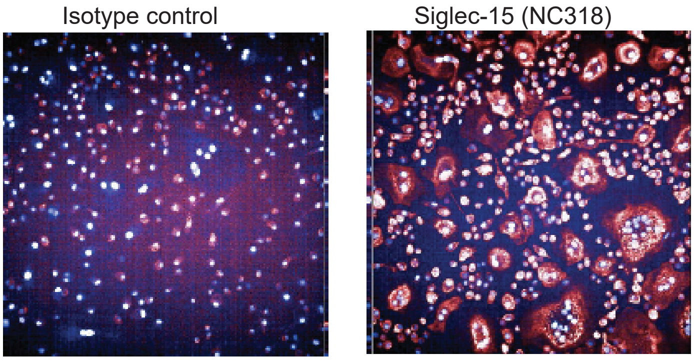


Primary human osteoclasts
